# Behavior decoding delineates seizure microfeatures and associated sudden death risks in mice

**DOI:** 10.1101/2025.01.13.632867

**Authors:** Yuyan Shen, Jaden Thomas, Xianhui Chen, Jaden Zelidon, Abigayle Hahn, Ping Zhang, Aaron Sathyanesan, Bin Gu

## Abstract

Behavior and motor manifestations are distinctive yet often overlooked features of epileptic seizures. Seizures can result in transient disruptions in motor control, often organized into specific behavioral patterns that can inform seizure types, onset zones, and outcomes. However, refined analysis of behaviors in epilepsy remains challenging in both clinical and preclinical settings. Current manual video inspection approaches are subjective, time-consuming, and often focus on gross and ambiguous descriptions of seizure behaviors, overlooking much of the intricate behavioral dynamics and action kinematics. Here, we utilized two machine learning-aided tools, *DeepLabCut* (DLC) and *Behavior Segmentation of Open Field in DLC* (B-SOiD), to decode previously underexplored behavior and action domains of epilepsy. We identified 63 interpretable behavior groups during seizures in a population of 32 genetically diverse mouse strains. Analysis of these behavior groups demonstrates significant differential behavior expression and complexity that can delineate distinct seizure states, unravel intrinsic seizure progression over time, and inform mouse strain backgrounds and genotypes. We also identified seizure behavior patterns and action/subaction kinematics that determine the risks of sudden unexpected death in epilepsy (SUDEP). These findings underscore the significant potential for translation into inpatient settings for video analysis in epilepsy monitoring units and outpatient settings via home surveillance devices and smartphones.

**One Sentence Summary:** AI-aided behavior decoding delineates intricate seizure microfeatures in mice.

## Introduction

Semiology constitutes a pivotal yet frequently overlooked aspect of epilepsy (*1*). Indeed, behavior and motion components of epileptic seizures are essential for the diagnosis of epilepsy, even with the advent of electroencephalography (EEG) and neuroimaging techniques (*2, 3*). Pleomorphic seizure behaviors also inform seizure types (*4*) and epileptogenic zones(*5*) and, therefore, guide the treatment. However, refined analysis of behavior in epilepsy is complex and cumbersome in both preclinical and clinical settings. Current studies focus on descriptive gross behaviors but discard the intricate transition dynamics and the richness of behavioral repertoire. Videography offers a simple method for recording behaviors in both epilepsy patients and animal models of seizures. Video monitoring is a routine practice in the epilepsy monitoring unit (EMU) to facilitate diagnosis. Ambulatory videos also provide predictive and additive value for diagnosing epileptic seizures in outpatients (*6-9*), particularly in telemedicine settings (*10, 11*). Additionally, video recordings serve as indispensable tools for preclinical studies using varying model organisms (*12*). However, extracting specific aspects of behavior for further analysis is ambiguous, subjective, nonquantitative, and highly time-consuming. The underinvestigated behavior domain harbors information vital for characterizing seizures and predicting their related outcomes.

The rapid advancements in artificial intelligence (AI) have revolutionized data-driven visual inspection, enabling fast, automated, and robust analysis of behavior and motion without user bias. Open-source tools for pose estimation and behavior classification have made fine motor movement tracking and prediction possible (*13*). For example, recent developments have highlighted the utility of AI-aided behavior analysis in studying epilepsy clinically (*14-16*) and preclinically (*17*). Work from *Gschwind et al*. employing 3D video recording and Motion Sequencing (MoSeq) has shed light on behavioral phenotypes in epileptic mice by tracking their naturalistic behavior during epileptogenesis (*18*). Despite these advances, significant knowledge gaps and technical barriers persist, particularly in uncovering finer behavior patterns and motifs, including action and subaction kinematics, and understanding how these findings are generalized across genetic backgrounds. Furthermore, The utility of seizure behavior in distinguishing and predicting yet indiscernible seizure outcomes, like seizure progression over time and sudden unexpected death in epilepsy (SUDEP), has not yet been demonstrated. Technological limitations also present challenges. For instance, MoSeq requires specialized 3D video recording setup and high storage and computational demands for processing large 3D video datasets, restricting its scalability and broader application (*19*). To overcome these barriers, we utilized two AI-aided tools, *DeepLabCut* (DLC) (*20*) and *Behavioral Segmentation of Open Field in DLC* (B-SOiD) (*21*), to analyze and integrate previously underexplored behavior domains of epilepsy based on 2D videos recorded using a single off-the-shelf camera in an entirely data-driven manner.

By leveraging DLC and B-SOiD, we focused on induced seizure behavior in a genetically diverse population of Collaborative Cross mice, with an emphasis on their motor motifs, behavior complexity, behavior patterns, and action/subaction kinematics in delineating seizure states, seizure propagation, and SUDEP. We also unraveled seizure phenotypes that are challenging to recapitulate in Angelman syndrome model mice with relevant pathogenic mutations. We identified serial behavior microfeatures that can be used to delineate seizure outcomes and predict seizure-related mortality in mice. This work highlights the transformative potential of integrating AI-aided tools with accessible video recording setups to achieve automated measures of action and subaction on a sub-second resolution, enabling high-throughput analysis of behavior at scales and significantly reducing the time and effort needed for manual scoring and analysis. Our work also paves the way for high-throughput and quantitative behavioral studies in both preclinical and clinical epilepsy research.

## Results

### Automated pose estimation and behavior group clustering using DLC and B-SOiD

We first trained DLC by manually labeling 28 body parts using a total of 240 frames of videos captured by a single side view camera from 12 mice covering all three coat colors commonly found in laboratory mice (i.e., albino, agouti, and black). We performed three rounds of refinement to achieve the benchmark of test error = 2.23 pixels (p-cutoff = 0.6). We then inferred the rest of the videos to generate pose estimations with the coordinates of each body part at 30 frames per minute across the preictal, myoclonic seizure (MS), and generalized seizure (GS) states of each mouse. We finally fed DLC pose estimations to B-SOiD unsupervised clustering using 30 mice, generating 67 unique behavior groups (BGs) (Fig. 1). Our B-SOiD model achieved 88% median accuracy on 20% of testing data, with the majority BGs (94.0%) reaching ≥80% accuracy (Suppl Fig. 1). We excluded four BGs (i.e., BG#53, BG#54, BG#57, and BG#59) from downstream analysis due to low clustering accuracy (< 80%) and percent usage (< 1%). The remaining 63 BGs were interpreted using general mouse ethogram and seizure-specific semiology (*1*) based on at least five representative snippets extracted for each BG (*21*) (Suppl Table 2).

**Figure 1.**
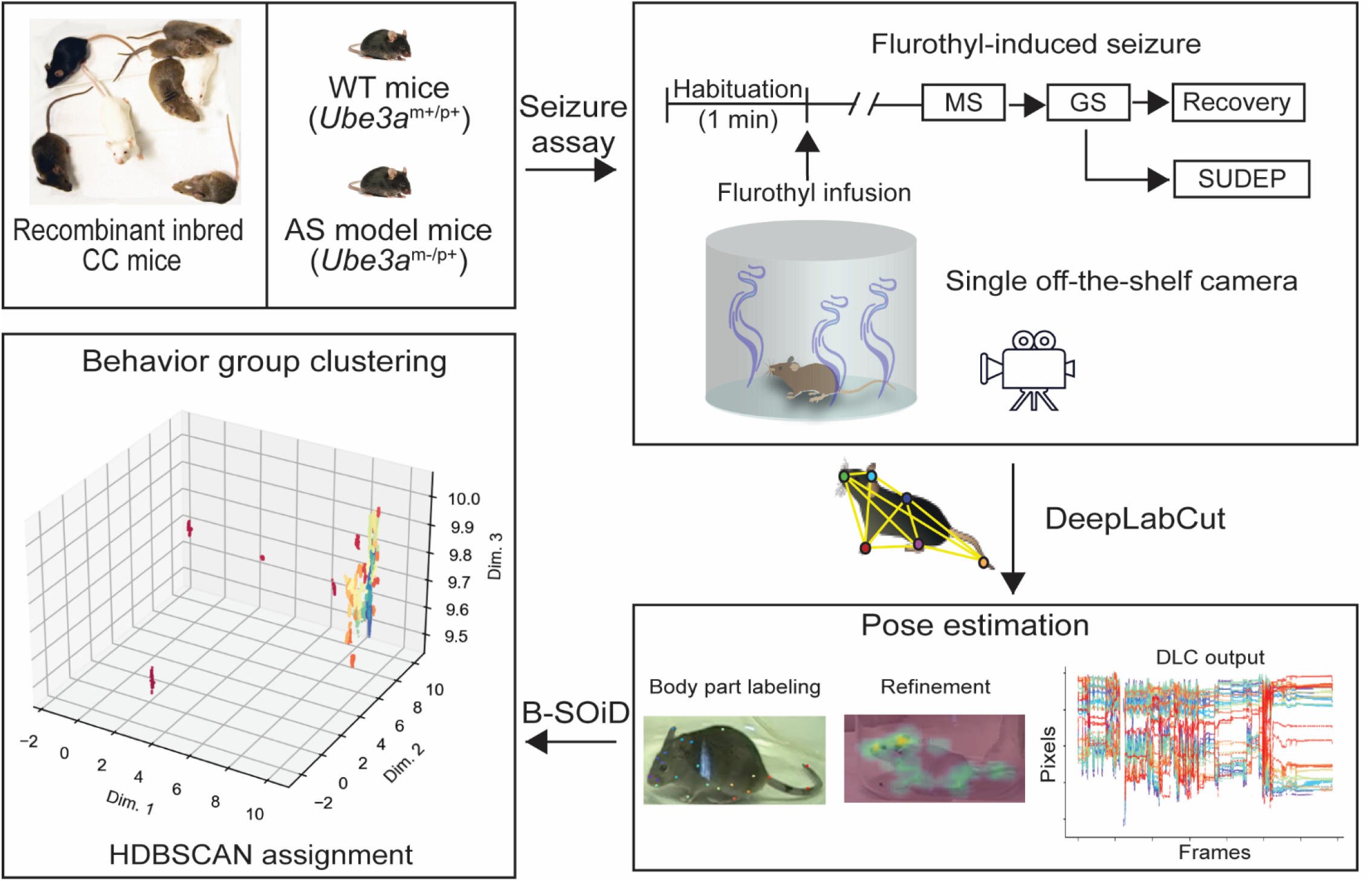
Schematic workflow depicts seizure assay using flurothyl challenge, 28 body parts pose estimation using DLC, and behavior groups (BGs) clustering using B-SOiD. Our model generates 67 BGs, 63 of which achieve ≥80% accuracy.

### Fine behavior discrimination of seizure states

Flurothyl is a commonly used chemoconvulsant that faithfully elicits evolving behavior seizure manifestations in mice several minutes after exposure. The state of flurothyl-induced seizure was traditionally classified manually into MS (i.e., intermittent jerks of the head and neck musculature) and GS (i.e., sustained body clonus with secondary loss of postural control) states based on visual inspection of the gross motor signs (*22*). However, Manual seizure state identification and characterization are subjective and time-consuming. We first tested whether our DLC/B-SOiD pipeline can extract the fine motor motifs and their dynamics that can be used to discriminate seizure states as a benchmark. Our first dataset consists of 219 mice (147 males and 72 females) from 31 inbred strains of Collaborative Cross mice in addition to the classic laboratory inbred C57BL/6J (C57) mice as a reference strain to capture diverse genetic backgrounds. We found the percent usage of 40 out of 63 BGs are significantly different (P < 0.05) across preictal, MS, and GS states (Fig. 2A). Among them, BG#40 (5.36% absolute and 115.1% relative change), BG#58 (3.93% absolute and 108.1% relative change), and BG#63 (3.25% absolute and 47.3% relative change) are the top three most differentially used BGs (average of |delta| > 3%). We then conducted principal component analysis (PCA), which generated new independent variables as linear combinations (*23*). PCA revealed a single significant PC1 (P = 0.006, 10,000 permutation) representing 13.01% of the variance. PC1 reflects the between-group differences separating preictal vs. MS (P < 0.001) and preictal vs. GS (P < 0.001), but not MS vs. GS (P = 0.416); whereas PC2, though not significant, represents 9.10% of the variance and better reflects the differences between MS vs. GS (P < 0.001) (Fig. 2B – 2D). We also calculated the loading of each BG in contributing to PC1 and PC2 and identified BG#40 (loading = 0.845) and BG#34 (loading = -0.846) as the most strongly related variables to PC1 and PC2, respectively (Suppl Fig. 2). Beyond the analysis of percent BGs usage, we next assessed the amplitude of randomness and complexity of behavior by computing the Shannon entropy (*24*). We found that entropy was significantly reduced during MS (P < 0.001) and GS (P < 0.001) compared to preictal state, suggesting that mice experienced more stereotyped behaviors after seizure onset regardless of seizure states (P > 0.99, MS vs. GS) (Fig. 2E and 2F). The sex dependency of innate and seizure behaviors remains under debate (*25-27*), therefore, we evaluated the sex effects on behavior compositions across preictal, MS, and GS states at the population level over 32 mouse strains. Surprisingly, males and females shared similar preictal behavior in terms of BGs usage, whereas they exhibited significant differential usage (P < 0.05) of specific BGs during MS (BG#11, BG#24, BG#51, BG#56, and BG#65) and GS (BG#34) (Suppl Fig. 3). This finding suggests sex-dependent seizure behaviors exist, emphasizing the inclusion of both sexes for seizure phenotyping.

**Figure 2.**
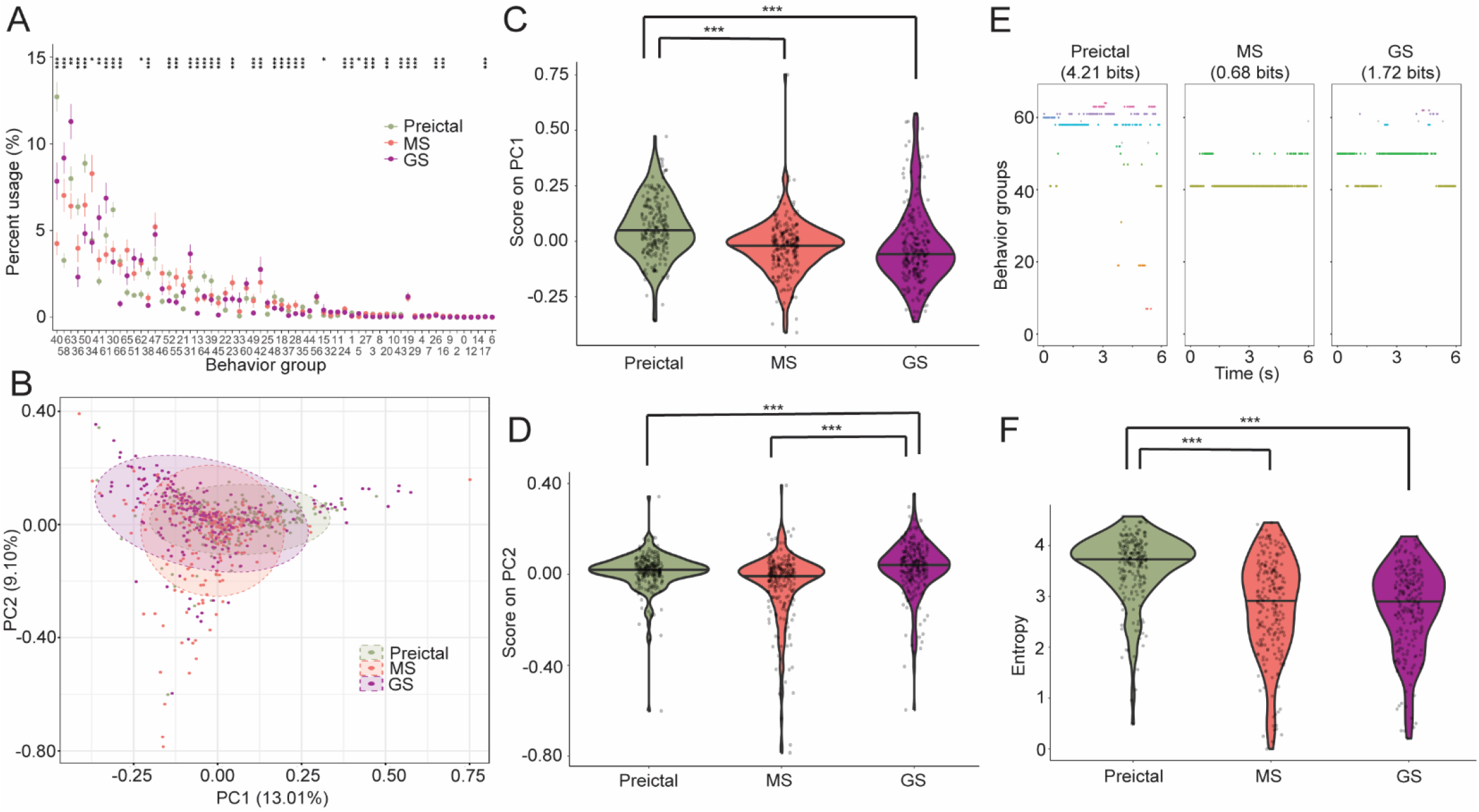
Distinct BG composition and complexity discriminate seizure states across strains. (A)Percent BGs distribution comparing preictal, MS, and GS ranked by the average of their absolute deltas in descending order. (B) PCA of BG usage with 95% confidence ellipse of (C) PC1 covering 13.01% of the variability and (D) PC2 covering 9.10% of the variability. (E) Representative BG usage of a mouse exhibited higher entropy (4.21 bits) during the preictal state compared to lower entropy during MS (0.68 bits) and GS (1.72 bits). (F) Analysis of behavior entropy by seizure states. Data were presented (A) as mean±SEM and analyzed using Friedman Test with Bonferroni adjustment; and (C, D, and F) as violin plots with median and individual data points and analyzed using Mann-Whitney U test. ^*^ P < 0.05, ^**^P < 0.01, and ^***^P < 0.001.

**Figure 3:**
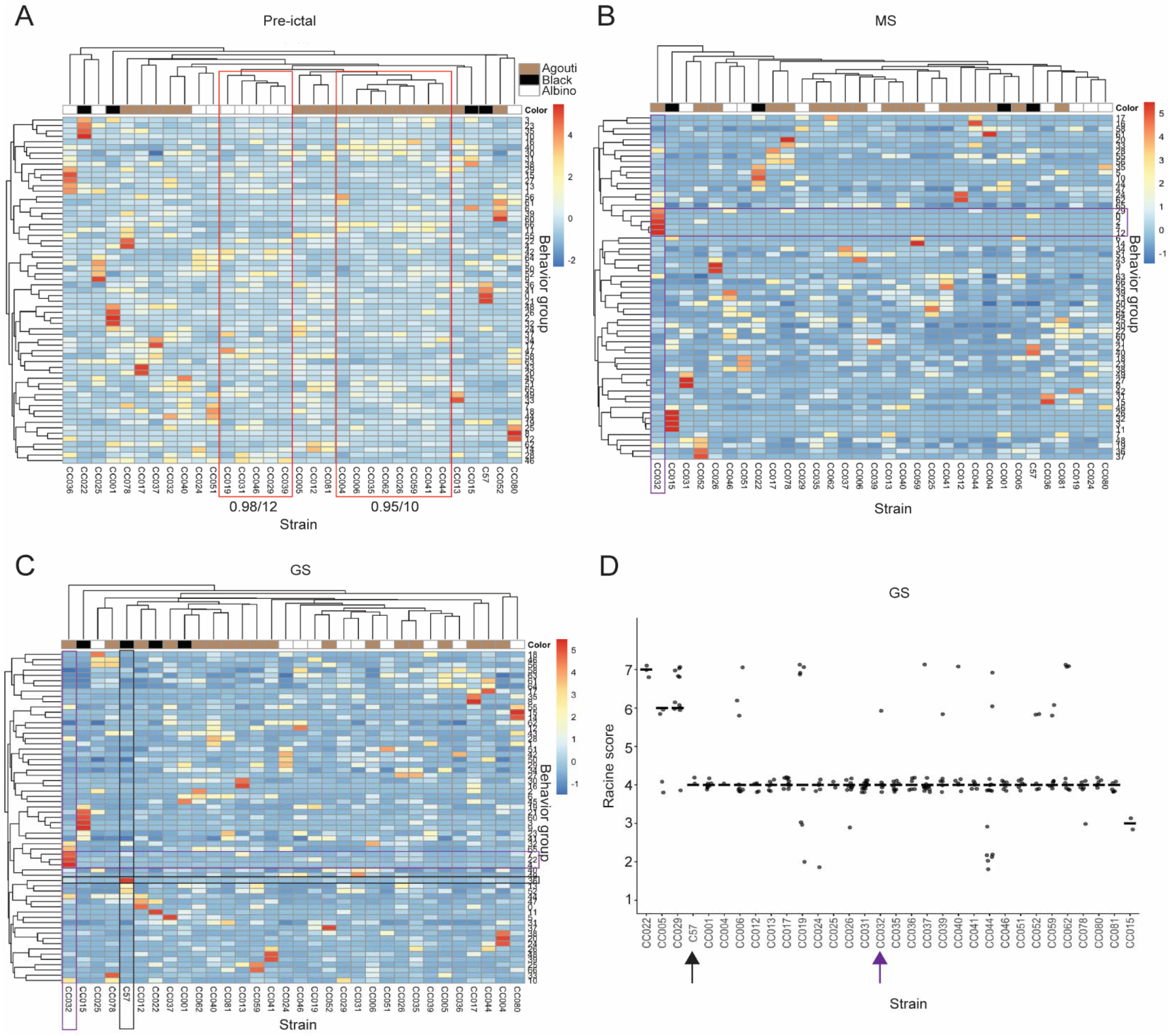
Hierarchical classification clusters distinct mouse strains based on BGs usage. Hierarchical clustering of the 32 inbred mouse strains, using percent usage of BGs during (A) preictal, (B)MS, and (C) GS. Each row represents a behavior group and each column represents a strain. Heatmap indicates standardized across strains percent usage above (red) and below (blue) the mean, respectively. Dendrograms of strains (above matrix) and behavior groups (to the left of matrix) represent overall similarities in BGs usage profiles and were clustered by similar values using complete linkage and an Euclidean method for distance. Red squares denote statistically significant clusters with AU P value ≥ 95% and number of bootstrap ≥ 10 labeled below. Purple and black squares denote the dominant (z score > 4) BG clusters in CC032 and C57, respectively, that are distinct from the rest of the mouse strains during (B) MS and (C) GS. (D) Racine scores of GS of each mouse with the median were analyzed using pairwise Mann-Whitney U test. Purple arrow denotes CC032 and black arrow denotes C57, which behavior is discernible by BGs classification but not Racine scores.

### Mouse strain genetic backgrounds dependent behaviors

Mouse genetic makeup largely determines their responses to seizure stimuli (*28, 29*), including those induced by flurothyl (*30, 31*). To reveal the mouse strain relationships based on their behavior response to seizure stimuli, we conducted a hierarchical clustering analysis using BG usage during preictal, MS, and GS across 31 CC and C57 strains. Clusters of mouse strains emerged, indicating correlated networks of these BGs usage patterns within each cluster (Fig. 3 and Suppl Fig. 4). Particularly remarkable, the most significant strain clusters (alpha ≥ 95% and bootstrap > 10) of preictal behavior were also associated with agouti (8 out of 8) and albino (5 out of 5) coat colors (Fig. 3A and Suppl Fig. 4). The agouti cluster exhibit more active movements, such as rearing (BG#66, difference in z score = 2.88) comparing to the albino cluster, who are relatively sedentary. These associated networks of behaviors indicate that mice with different coat colors may follow a neurodevelopmental trajectory that can be broadly characterized using fine behavioral networks. Indeed, coat color has long been associated with behavior (*32, 33*). A genotype-based analysis suggested that agouti and albino loci affect exploratory activity (*34*). Of note, we trained our DLC model using a balanced number of albino, agouti, and black mice to minimize the coat color bias during model training.

**Figure 4.**
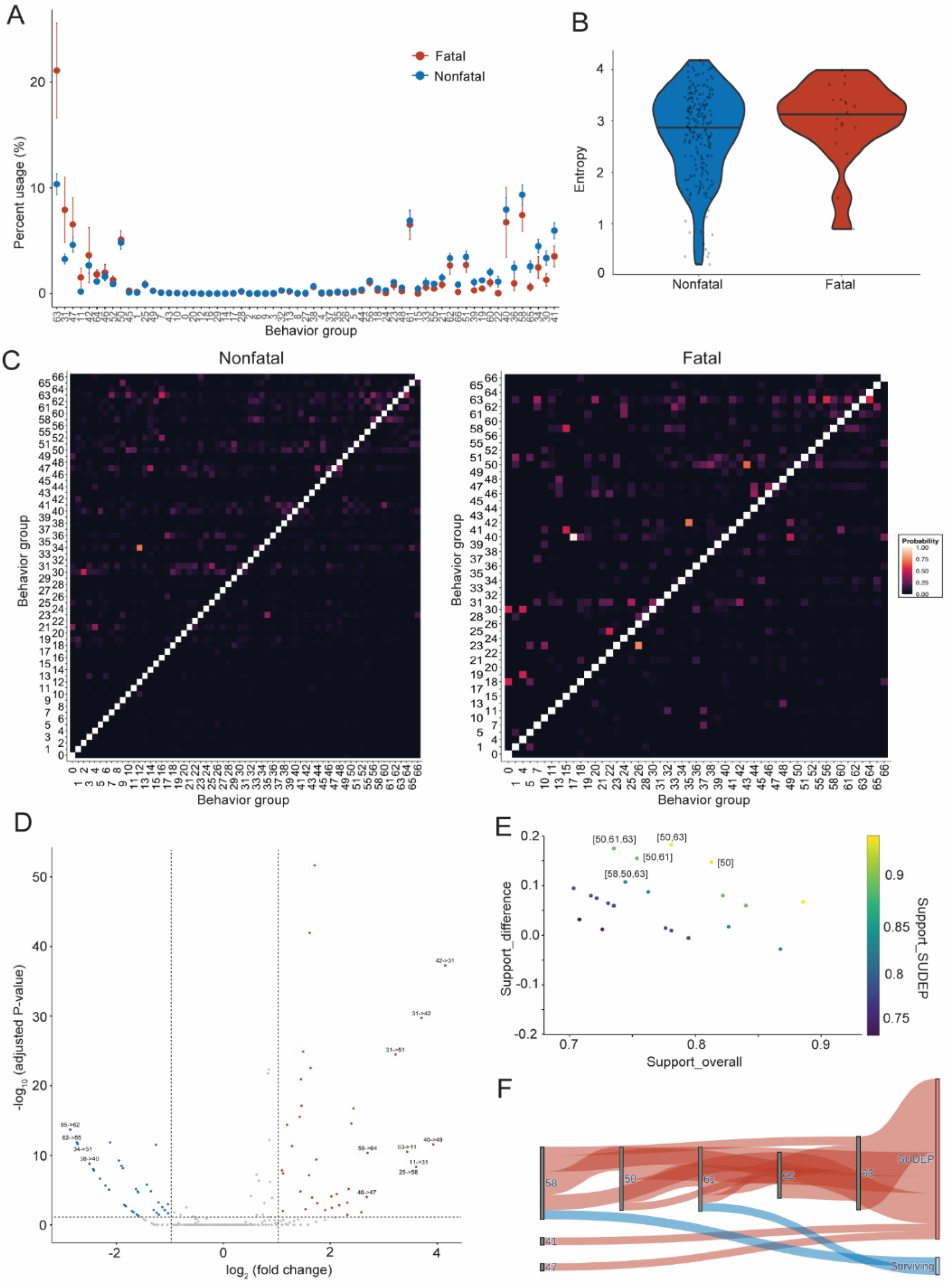
Behavior microfeatures, transitions, and frequent pattern mining discriminate the mortality of seizures. (A) Percent BGs distribution comparing fatal and nonfatal seizures during GS, ranked by the delta of fatal and nonfatal seizures in descending order. Data are presented as mean ± SEM and analyzed using the Mann-Whitney U test. (B) Behavior entropy during fatal and nonfatal seizures was presented using violin plots with median and individual data points and analyzed using Mann-Whitney U test. (C) Probability transition matrix of nonfatal and fatal seizures. Particular BGs that were not utilized during GS of fatal seizures were excluded from plotting. (D) Volcano plot represents differential expression of BGs transition during fatal compared to nonfatal seizures with a threshold of adjusted P value < 0.05 and |log2 fold-change| > 1. Data points reached a more stringent adjusted P value <0.01 and |log2 fold-change| > 2 were labeled. (E) The scatter plot visualizes the relationship between the support of itemsets and the difference in support between fatal and nonfatal seizures with ≥ 70% support overall and ≥ 50% support within each group. The top five itemsets with the largest different support were labeled. (F) Sankey diagram illustrates the most frequently occurring itemsets comparing fatal seizures leading to SUDEP with nonfatal seizures where the mouse survived.

Although no statistically significant clusters were identified during MS or GS (Suppl Fig. 4), different CC strains displayed pleomorphic seizure behaviors (Fig. 3B and 3C). For example, we found that CC032 exhibited a unique behavioral cluster during MS (BG#29, BG#0, BG#2, BG#4, and BG#12) and GS (BG#7, BG#22, and BG#4) characterized by a collection of dominant BGs distinct from other mouse strains (z score > 4). The commonly used laboratory inbred C57 also demonstrated distinctive behavior representations of freezing with forelimb stretching (BG#36, z score = 5.27) during GS. However, these behavior signatures were not discernible during GS if analyzed using the traditional semi-quantitative Racine score (P = 1.0) (Fig. 3C and 3D). These observations support the notion that the epileptic behavior phenotype obtained from a particular mouse strain, including the commonly used C57 in epilepsy studies, may not be generalizable across diverse genetic backgrounds.

### Behavior biomarker of SUDEP

SUDEP is the most devastating consequence of seizures and lacks prognostic biomarkers (*35*). A recent video analysis suggested that abnormal movement during convulsions leads to SUDEP in limited cases under home video surveillance (*36*). We thereby first focused on the usage of BGs and their entropy during the early clonic phase of GS. Notably, the percentage of most differentially utilized BG#63 doubled during fatal (21.10%) compared to nonfatal seizures (10.53%). However, this difference did not reach statistical significance after adjustment of multiple comparisons (P = 0.076) (Fig. 4A). The behavior complexity measured by entropy was also similar between fatal and nonfatal seizures (P = 0.168) (Fig. 4B). Albeit similar static BGs composition, we assessed the prevalence of transition of two adjacent BGs and found 13 BGs transitions were differentially expressed during fatal comparing to nonfatal seizures that passed the stringent threshold of adjusted P value < 0.01 and |fold-change| > 2. The transitions probability of BGs 42→31, 31→42, 31→51, 40→49, 63→11,58→64, 11→31, 25→58, and 40→47 were increased and transitions probability of BGs 55→62, 62→55, 34→51, and 38→40 were decreased during fatal compared to nonfatal seizures (Fig. 4C and 4D). This result suggests a dynamic reorganization of ictal motor activity that may reflect the involvement of differential circuit mechanisms underlying fatal outcomes (*37, 38*).

To further identify recurring patterns, relationships, and dependencies among different BGs for SUDEP in a data-driven manner, we employed an unsupervised Apriori machine learning algorithm for frequent pattern mining and association rules learning to uncover hidden patterns (i.e., itemsets) between classes of “fatal” and “nonfatal” seizures. We focused on the itemsets that not only occur preferentially in one class over the other but also with high frequency (≥ 70% overall support and ≥ 50% for each class), ensuring they serve as strong indicators of their respective classes (Fig. 4E). A Sankey diagram further illustrated the most commonly used itemsets for each class and revealed particular connections cross BG#58 (diving), BG#50 (dropping), BG#61 (resting and gasping), BG#62 (clonus with straub tail), and BG#63 (balancing) in segregating fatal seizures in contrast to nonfatal seizures. The Sankey diagram of frequent patterns also underscored the effectiveness and sensitivity of the frequent pattern mining, revealing a higher probability for fatal outcomes despite their number (n = 19) being ten times less than that of surviving mice (n = 200) (Fig. 4F). We finally employed association rules analysis by measuring lift of distinct collections of strongly associated itemsets between fatal and nonfatal seizures. We identified the top ten most useful itemset associations for each class to facilitate more intuitive and faster classification in the real world (Suppl Fig. 5).

In addition to BGs composition and transition, we further examined the kinematics of particular motions during GS that are associated with SUDEP. We first focused on the cumulative distribution function (CDF) of BG bout duration and found a profound left shift of BG#23 (pushing backward, P = 0.045) and right shift of BG#31 (leaping, P = 0.005), with fatal seizures displaying shorter pushing and longer leaping motions compared to nonfatal seizures (Fig. 5A and Suppl Fig. 6). Converging lines of evidence also suggest that mice typically experience tonic seizures characterized by sudden muscular stiffness in extremities that may involve the brainstem and finally lead to SUDEP (*39*).

**Figure 5:**
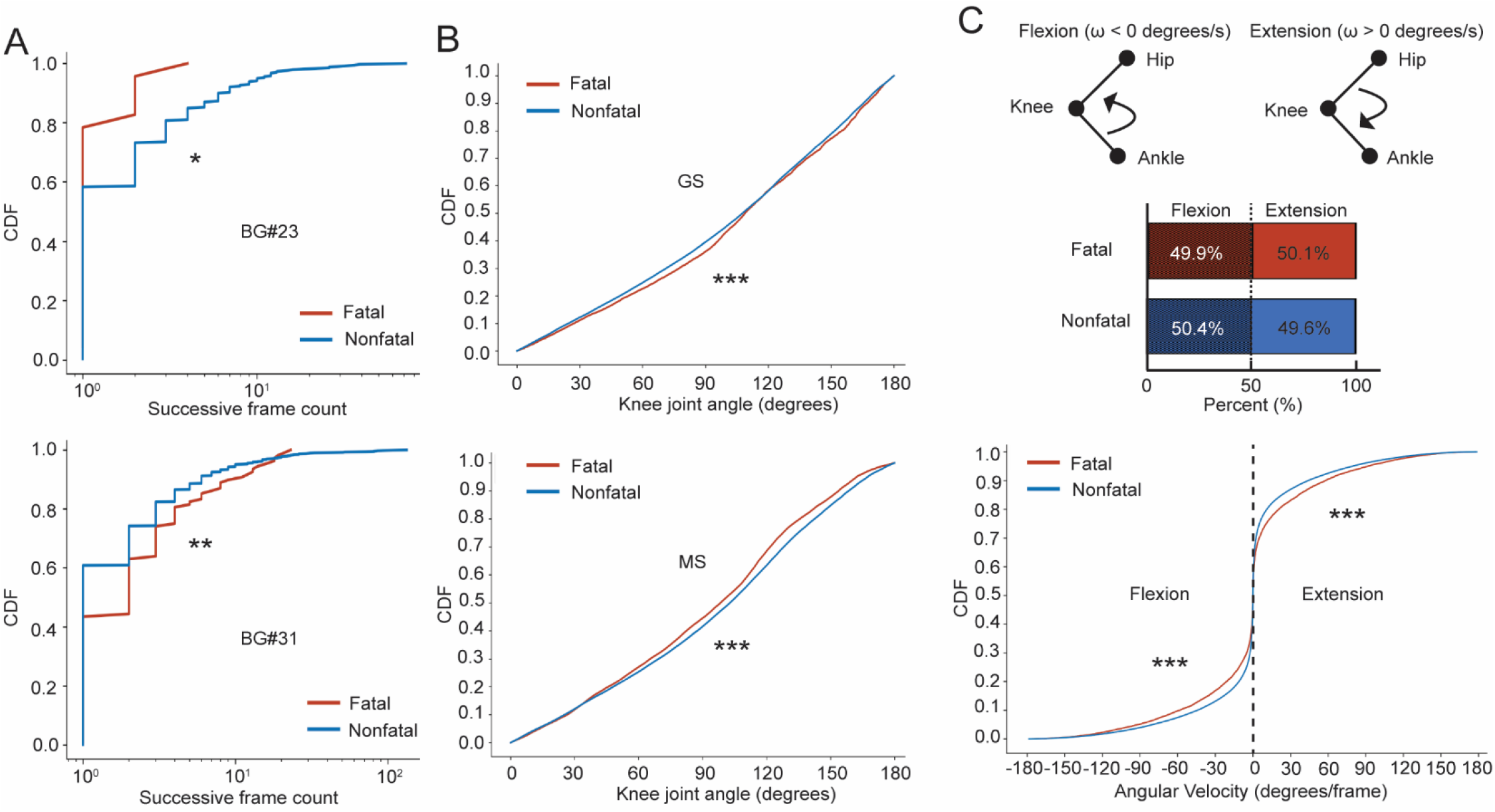
Ictal behavior and motion kinematics determine mortality of seizures. (A) CDF of bout duration of BG#23 and BG#31 during GS of fatal vs nonfatal seizures. (B) CDF of knee joint angles during GS of fatal vs nonfatal seizures. (C) Direction and speed of knee joint motion measured by angular velocity (flexion: ω < 0 degrees/s and extension ω > 0 degrees/s) during GS of fatal vs nonfatal seizures. CDF was analyzed using two-tailed Kolmogorov-Smirnov test, ^*^P□<□0.05, ^**^P□ < □ 0.01, and ^***^P□ < 0.001.

However, tonic seizure is not a valid behavior indicator of SUDEP, as mice can survive after experiencing tonic seizures (*40, 41*). Instead of gross classification of the presence or absence of tonic hindlimb extension, we focused on the knee joint angles during the clonus phase of GS to quantitatively assess the correlation between the ictal hindlimbs posture and mortality. Knee joint angle CDF suggests mice displayed increased knee joint angles during fatal compared to nonfatal GS, a shift from MS states (Fig. 5B). Finally, we calculated an instantaneous angular velocity, which allows us to determine the turning direction and speed of knee joint manifested to seizures. We first focused on the direction of knee motion and found comparable distribution of flexion and extension during fatal and nonfatal seizures (Fig. 5C). However, knee joint angular velocity CDF suggests faster hindlimb flexion (P < 0.001) and extension (P < 0.001) during fatal vs nonfatal seizures (Fig. 5C).

These data suggest motion kinematic measurements, including bout duration, as well as knee joint angles and speed, can further segregate seizure mortality.

### Seizure behavior evolves over time during repeated seizure challenges

Kindling models the process of epileptogenesis with the concept that seizures progress over time or repeated seizures render the subject more susceptible to subsequent seizures (*42*). For example, the seizure threshold was reduced in most commonly used laboratory inbred mice strains over repeated flurothyl exposure, suggesting elevation of seizure sensitivity after exposure to repeated seizures (*22*). However, the evolution of behavior seizure severity and characteristics over time remains contradictory. Some studies suggest seizure behavior evolves over time (*43*), while others argue the behavioral components of epileptic and functional seizures are usually stereotypic and repetitive from one seizure episode to another for a particular subject (*44-46*). We previously identified CC051 mice as a kindling-resistant CC strain, with their seizure threshold remaining unchanged across 8-day flurothyl kindling (*31*). Despite similar behavior seizure thresholds and gross assessments of seizure severity, we hypothesize that the microfeatures and complexity of their behavior seizures evolve over repeated episodes. We video-recorded a separate cohort of CC051 mice and confirmed their MST, GST, and Racine scores remain similar across 8 days of flurothyl kindling (Fig. 6A). We subsequently examined the daily dynamics of their behavior components usage and complexity over time. 21 and 19 out of 63 BGs exhibited above the average percent usage difference on day 8 compared to day 1 during MS and GS, respectively (Fig. 6B and 6C). Among them, we found the percentage usage of BG#47 (freezing with minor head shaking, P = 0.023) during MS and BG#40 (scrambling up, P = 0.023) during GS was significantly increased over time (Fig. 6B and 6C). Behavior complexity analysis also revealed a significant shift of entropy during GS across 8 days (P = 0.042) with the seizure behavior complexity peaking at day 5 (P = 0.012), suggesting a transient adaptation of motor response to seizure stimuli before it is stabilized (Fig. 6D). Interestingly, the preictal behavior complexity remains relatively consistent with a moderate decreasing trend over 8 days, suggesting the mice gradually acclimate to the testing camber before seizure onset (Fig. 6D). These results together suggest that behavior decoding unveils hidden temporal behavioral evolution over repeated seizures.

**Figure 6.**
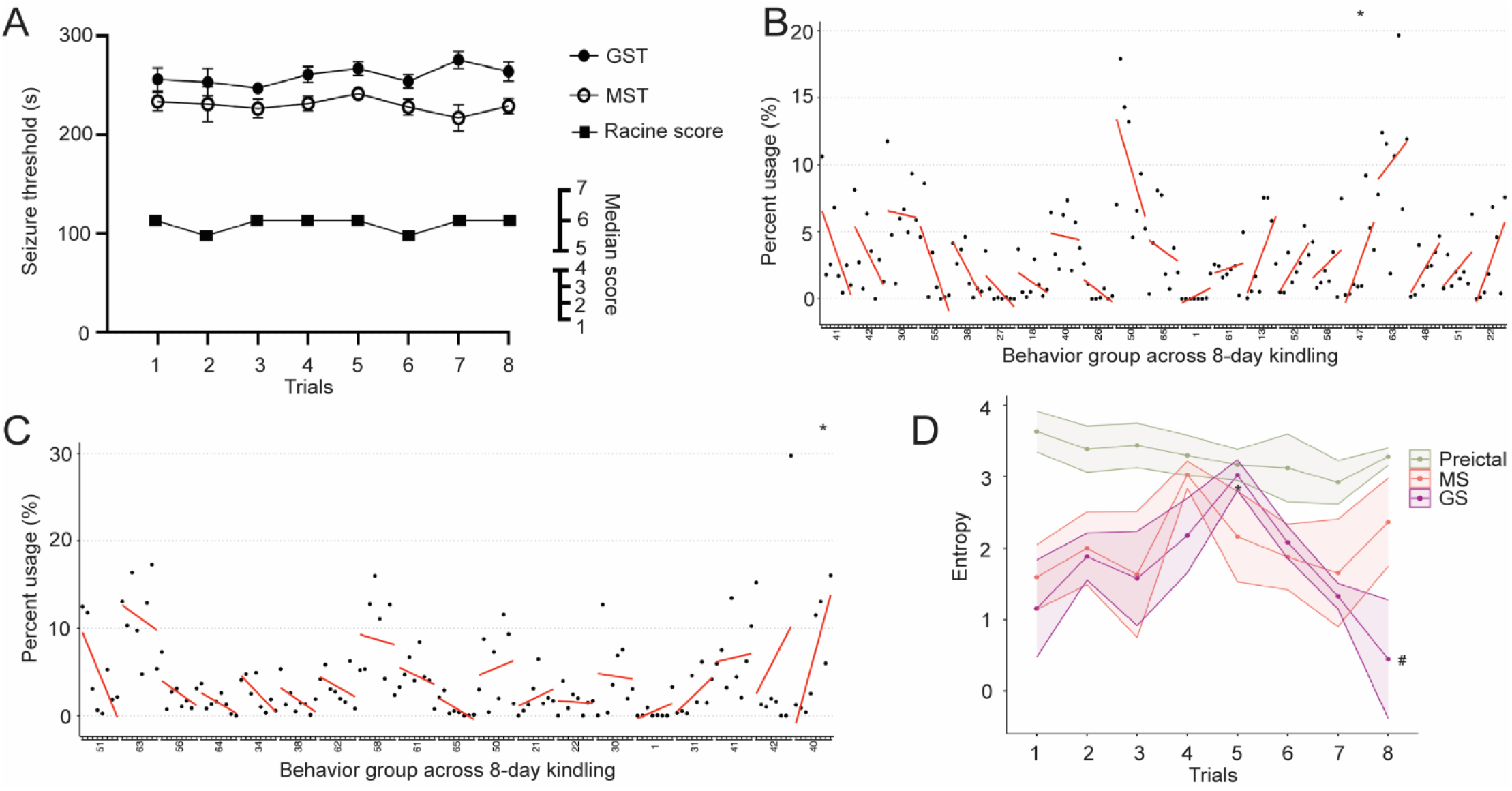
Seizure behavior evolves during flurothyl kindling over time. (A) similar MST, GST, and Racine scores of CC051 mice across 8 trials of flurothyl kindling. (B-C) Percent usage progression over time across 8 days (minor ticks on the x-axis within each BG) of BGs above average absolute delta between day 8 and day 1 in ascending order based on the slope of linear regression line during (B) MS and (C) GS. Data were presented as mean daily usage and analyzed using a linear mixed effects model with Bonferroni test, ^*^P < 0.05. (D) Summary of behavior entropy during preictal, MS, and GS across 8 days of kindling. Data were presented as mean±SEM and analyzed using ANOVA, ^#^P < 0.05, and linear mixed effects model, ^*^P < 0.05.

### Fine seizure behavior phenotyping of Angelman syndrome model mice

Transgenic mouse models have been commonly used to study particular pathogenic mutations in contributing epileptic phenotypes by measuring their responses to seizure stimuli. However, many epileptic phenotypes can not be recapitulated in their corresponding transgenic animal models or in a strain background-dependent manner (*47*). For example, mutation or deletion of the *UBE3A* gene results in Angelman syndrome, with over 80% of patients developing epileptic seizures within 3 years of age (*48*). However, mice lacking the *Ube3a* gene, on C57 background, did not exhibit typical spontaneous seizure and showed similar seizure susceptibility to their initial exposure to flurothyl as measured by MST, GST, or Racine score compared to WT littermates (*49*) (Fig. 7A-7C). We tested whether the quantitative behavior analysis could reveal distinct behavior microfeatures that can discriminate WT and AS model mice, despite their similar seizure thresholds and Racine scores. We video-recorded flurothyl-induced seizure in naïve WT and AS mice and subjected their video recordings of MS and GS to DLC/B-SOiD. We found BG#34 (10.68% absolute and 67.2% relative decrease) during MS and BG#62 (9.79% absolute and 67.7% relative decrease) during GS have the largest percent usage change in AS mice compared to WT. However, after adjusting for multiple comparisons, they do not reach statistical significance (Fig. 7D and 7F). Notably, AS model mice displayed reduced ictal behavioral seizure stereotypies, with their entropy significantly higher compared to WT during GS (P < 0.01). In contrast, their entropy levels were comparable during MS (P = 0.70) (Fig. 7E and 7G). Collectively, these results reveal previously unrecognized behavior microfeatures that can discriminate seizure phenotype in AS mice, which is indiscernible to traditional descriptive or semi-quantitative behavior analysis.

**Figure 7.**
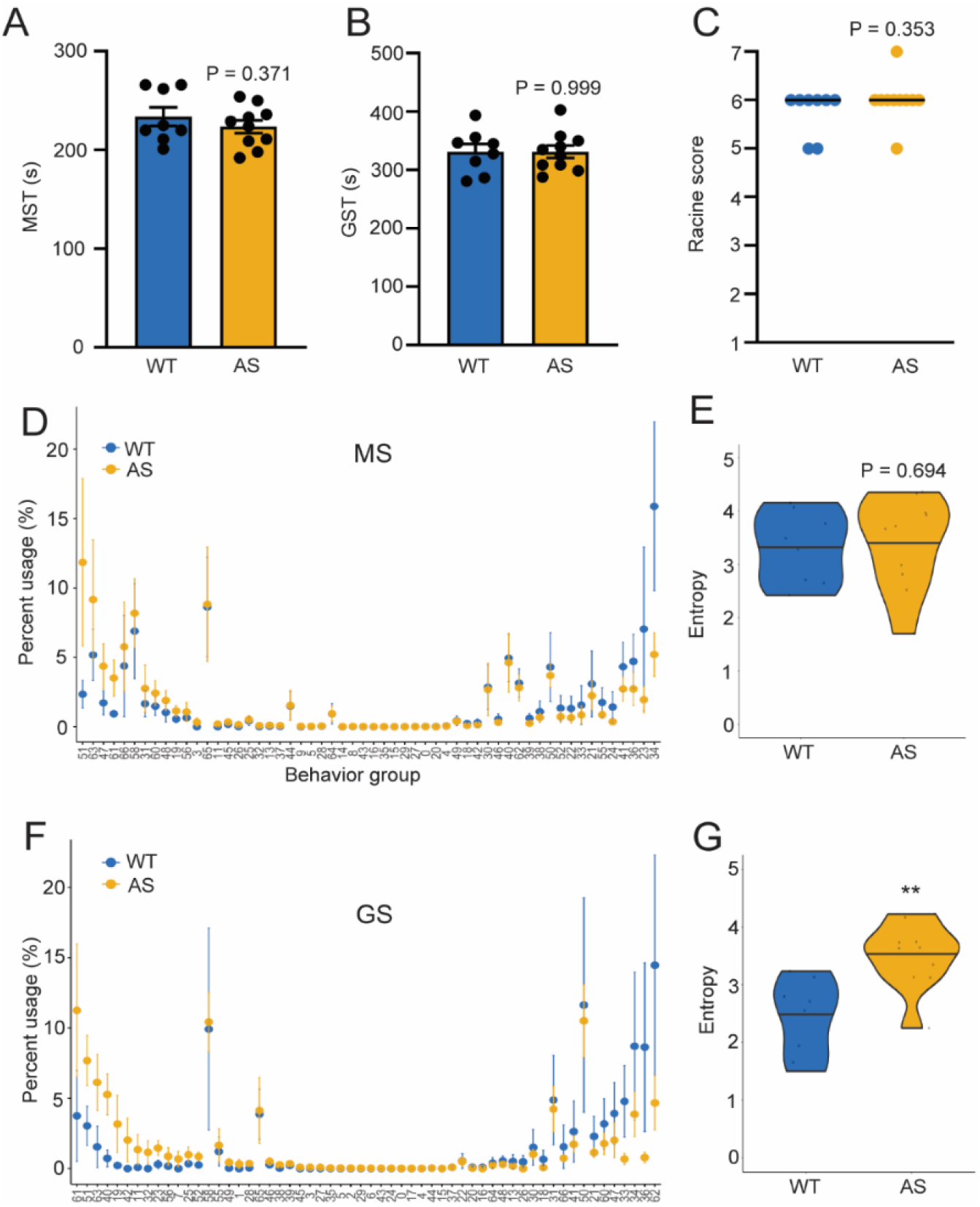
AS model mice exhibit higher generalized seizure complexity. (A) MST, (B) GST, and (C) Racine score of single flurothyl-induced seizure in WT (n = 8) and AS (n = 10) model mice. (D) and (F) Percent BGs distribution comparing WT and AS model mice during (D) MS and (F) GS ranked by the delta of AS and WT mice in descending order. (E) and (G) Behavior complexity measured by entropy during (E) MS and (G) GS of WT and AS model mice. Data is presented as (A, B, D, and F) mean±SEM and analyzed using (A and B) Student’s t-test and (D and F) Wilcox Test with Bonferroni adjustment; as (C) median plus data points and (E and G) violin plots with median and individual data points and analyzed using Mann-Whitney U test, ^**^P < 0.01.

## Discussion

We employed the DLC/B-SOiD behavior decoding pipeline to analyze videographic recordings of seizure behavior in a genetically diverse population of recombinant inbred CC mice and a transgenic mouse model that harbors maternal deletion of *Ube3a* recapitulating Angelman syndrome. We profiled the behavior microfeatures that can delineate seizure states, progression, and mortality, as well as infer mouse strains and genotypes. Our study proved the utility of DLC/B-SOiD based behavior analysis in multiple domains of behavioral seizure study with greater precision. The minimal requirement of the experimental setup (single off-the-shelf camera) and superior tolerance to recording variation of DLC (e.g., light, distance, angle, background, etc.) allow for broader implications of this pipeline and secondary analysis of existing mice videographic datasets without additional training. Our approach reduces the time and financial resources traditionally required for manual video reviewing and behavioral analysis, ensuring reproducibility across laboratories for preclinical studies. Moreover, both DLC and B-SOiD are accessible to biologists without extensive coding knowledge or computational resources like high-performance GPUs. Ultimately, DLC/B-SOiD will empower researchers to conduct high-throughput and quantitative behavior seizure studies, accelerating discoveries in epilepsy and related disciplines. Our study also provides a foundation and opens up possibilities of implementing other available AI pose estimation (e.g., SLEAP, AniPose, MARS, etc.) and behavior clustering (SimBA, Keypoint-MoSeq, DeepEthogram, TREBA, etc.) tools to epilepsy studies (*13*).

Our study promises great translational potential into other model organisms and, finally, epilepsy patients. DLC has been successfully applied to studying behaviors across morphologically diverse species and scenarios (*50*), allowing adaptation of our pipeline in mouse models to include other model organisms, such as fruit flies, zebrafish, rats, and pigs, which have been frequently used in studying epilepsy (*51*). It is worth noting that DLC equipped with Model Zoo SuperAnimals offers pre-trained pose estimation models that can be used on over 45 species without additional human labels (*52*). Technically, both DLC and B-SOiD have been successfully employed for human pose estimation and behavior clustering. Videographic recordings are also integral parts of ICU, EMU, and ambulatory monitoring (*53*), warrant extensive and ever-increasing inpatient and outpatient videographic datasets available for studying naturalistic and epileptic behavior in humans. The automated behavior analysis is particularly valuable for videos acquired in the context of telehealth using home surveillance (*9*) and ubiquitous smartphones (*6, 7*). Therefore, there is a clear path to move quickly and efficiently from preclinical to clinical populations to improving video recordings as an adjunctive tool in diagnosing seizure-like events and predicting their related outcomes.

Our dataset of flurothyl behavior seizure assay in a population of recombinant inbred CC mice offers unique opportunities to explore how genetics govern behavior responses to seizure stimuli. The unprecedented extreme seizure outcomes, including increased susceptibility to SUDEP and resistance to epileptogenesis in a kindling model in particular CC strains, allow us to unravel the behavior components underlying these seizure-related outcomes that were previously challenging to study. For example, we identified a collection of intrinsic behavior microfeatures and dynamics that could discriminate and predict the mortality of preceding seizures and unravel seizure behavior characteristics that evolve over time. By employing DLC/B-SOiD pipeline in the CC mice population, we were able to establish a behavior repertoire for seizure responses considering the genetic complexity.

Our initial approach in identifying behavioral SUDEP indicators focused solely on the frequency of each BG without leveraging pattern mining. However, this approach resulted in insignificant indicators for identifying fatal seizures. Furthermore, in real-world applications, frequency counts can only be obtained after the entire generalized seizure episode has concluded, making it challenging to capture real-time changes and requiring post-event statistical analysis. To address these limitations, we employed a frequent pattern mining method, the Apriori algorithm, allowing us to move beyond static frequency counts and incorporate the relational context of BGs. This approach captures subtle yet significant and interpretable patterns, enabling the model to distinguish between classes more effectively.

The ability of quantitative seizure behavior decoding offers opportunities to elucidate the neural basis that regulates particular seizure-related behaviors. High temporal resolution (in milliseconds scales) behavior decoding of seizures in the time domain will enable the mapping of behavioral actions to neural activity using concurrent video-calcium/voltage imaging or -electrophysiology (e.g., single unit recordings) to understand the neuronal and circuit mechanisms underlying the behavior manifestations of epileptic seizures. This type of combination has already proven feasible with platforms like Consistent EmBeddings of high-dimensional Recordings using Auxiliary variables (CEBRA), an AI-driven technology capable of mapping behavioral actions to neural activity across species (*54*). Inferencing dense neural and behavioral data will be useful for revealing mechanistic relationships between the dynamics of neural activity and action and improving risk stratification of patients based on their endophenotypes, including behavior. Our study also opens up the possibility of integrating behavior features and motifs with neurophysiological and neuroimaging measures for more accurate and comprehensive multimodal seizure and SUDEP detection and prediction. Equally important, the capability of real-time pose estimation (within 15 ms, >100 FPS) of DLC (*55*) open the possibility of noninvasive behavior-triggered closed-loop seizure intervention. It is possible to use real-time pose estimation as a “trigger” for on-demand optogenetic and electrical stimulation to interrogate the underlying mechanisms of epilepsy and develop potential therapeutic interventions to halt seizures (*56*) and prevent SUDEP (*57*) when it is needed.

Some limitations and challenges remain that warrant further investigation. We utilized a single side-view camera, a common video-recording setup for monitoring spontaneous and induced seizures in lab animals. However, the side-view perspective requires more imputation in DLC labeling due to the occlusion of the body part facing away from the camera. Despite the occasional occlusion, a side-view camera offers advantages in visualizing detailed seizure-relevant extremity movement by capturing carpus, tarsus, knees, elbows, and tarsal joints that are commonly occluded from top views. It also better accommodates vertical actions like rearing, dropping, and straub tail. Climbing emerged as a prominent behavioral group in our study, possibly reflecting the limited space within the gas chambers (13 cm O.D. x 20 cm height cylinder) and the natural escape behavior of mice, which may be less frequent in open spaces. We also noticed that higher frame rate and resolution could facilitate more accurate labeling of body parts during the fast motor bouts seen in generalized tonic-clonic seizure activity during GS, such as jumping and “popcorning”. Although a sampling rate of 30 fps is typically sufficient for capturing sudden and rapid motion during seizures in people (*58*), mice generally exhibit more rapid motions than humans, suggesting that a higher frame rate would provide more precise body part prediction.

## MATERIALS AND METHODS

### Mice

Dataset 1 comprises 219 mice from 31 CC strains: CC001/Unc, CC004/TauUnc, CC005/TauUnc, CC006/TauUnc, CC012/GeniUnc, CC013/GeniUnc, CC015/Unc, CC017/Unc, CC019/TauUnc, CC022/GeniUnc, CC024/GeniUnc, CC025/GeniUnc, CC026/GeniUnc, CC029/Unc, CC031/GeniUnc, CC032/GeniUnc, CC035/Unc, CC036/Unc, CC037/TauUnc, CC039/Unc, CC040/TauUnc, CC041/TauUnc, CC044/Unc, CC046/Unc, CC051/TauUnc, CC052/GeniUnc, CC059/TauUnc, CC062/Unc, CC078/TauUnc, CC080/TauUnc, and CC081/Unc. We also studied classical laboratory inbred C57BL/6J (C57, JAX:000664) mice as a reference strain. The number, sex, age range, and coat color of each mouse strain were summarized (Suppl Table 1). A subset of dataset 1 was obtained from seizure phenotype screening for a genetic mapping study (*31*). Dataset 2 consists of 8-day flurothyl kindling data of 6 male CC051 mice. Dataset 3 consists of 10 (6 males and 4 females) Angelman syndrome model mice and 8 (4 males and 4 females) wildtype littermates (JAX #: 016590). All animal procedures were approved by the Institutional Animal Care and Use Committee of The Ohio State University and were performed in accordance with the guidelines of the U.S. National Institutes of Health. Mice were group-housed on a 12:12 light/dark cycle with *ad libitum* access to food and water.

### Flurothyl-induced seizure and video recordings

Each mouse was placed in a 2L glass beaker, and 10% flurothyl (bis-2,2,2-trifluoroethyl ether; Sigma-Aldrich) in 95% ethanol was then infused at a rate of 200 μL/min onto a disk of filter paper suspended at the top of the chamber. Mice exhibit various stages of increasing seizure severity in response to flurothyl exposure, including myoclonic seizure (MS, sudden involuntary jerk/shocklike movements involving the face and neck musculature) and generalized clonic seizure (GS, also known as clonic-forebrain seizures that are characterized by clonus of the face and limbs, loss of postural control, rearing, and falling). Upon emergence of a generalized seizure, the lid of the chamber was immediately removed, allowing for rapid dissipation of the flurothyl vapors. Most seizures resolved after the mouse was exposed to fresh air, whereas seizures in some mice progressed into the tonic phase with limb extensions that may lead to respiratory arrest and SUDEP. For flurothyl kindling, flurothyl exposures were repeated once daily over eight consecutive days. The mouse behavior during a flurothyl-induced seizure was recorded using a single off-the-shelf camera (JVC GZ-MG300 or SONY HDR-CX405 camcorders) from a side view at the resolution of 480 p and 30 frames per second. The videos were manually reviewed by investigators (blind to strains) who determined latency to the onset of myoclonic seizures and generalized seizures. For flurothyl kindling, flurothyl exposures were repeated once daily over eight consecutive days.

Behavior seizure was scored based on a modified Racine scale as described earlier (*22*). GS behaviors were classified using the following behavioral scoring system: grade 1, purely clonic seizure; grade 2, “transitional” behaviors involving high-frequency/low-magnitude bouncing and/or rapid backward motion; grade 3, running/bouncing episode; grade 4, secondary loss of posture with bilateral forelimb and hindlimb treading; grade 5, secondary loss of posture with bilateral forelimb tonic extension and bilateral hindlimb flexion followed by treading; grade 6, secondary loss of posture with bilateral forelimb and hindlimb tonic extension followed by treading; grade 7, death.

### DLC pose estimation

To train the DLC model, we used 20 image frames per mouse of 12 mice covering all three coat colors observed in this population (i.e., 4 black, 5 agouti, and 3 albino). We preprocessed each video by cropping out the chamber that constrains the mouse and trimming each cropped video into three time windows based on seizure states of preictal (2 mins prior to the onset of MS), MS (from the onset of MS to the onset of GS), and GS (from the onset of GS to either termination of convulsions or the emergence of tonic extension, whichever happens earlier). The model was pre-trained to detect and track 28 different body parts (snout, mouth, right eye, left eye, dorsal neck, ventral neck, right shoulder, left shoulder, right elbow, left elbow, right paw, left paw, dorsal body center, ventral body center, right hip, left hip, right ankle, left ankle, right center foot, left center foot, distal tail, center tail, and proximal tail) across the body. The model was further trained for up to 1,030,000 iterations using a deep residual network structure (ResNet-50) based on the pre-trained model weights from DLC. All DLC training and inferring were conducted.

### B-SOiD behavior clustering

To reduce the background noise in feature clustering, the original recordings were cropped in size (480 × 960) to only include the mouse within the edges of the glass container. These videos were then trimmed in length to include only the preictal (120 seconds immediately preceding MS), MS, and GS time windows using Shotcut. All cropped and trimmed videos were processed by the trained DLC model to obtain the body part positions for B-SOiD clustering. To predict the specific behaviors in a mouse by clustering, a B-SOiD model was created using the recording of preictal, MS, and GS time windows of 30 mice, allocating 80% for training and 20% for testing. All 28 body parts were used to extract features. The pose relationships of the body parts were first reduced into a lower dimensionality using Uniform Manifold Approximation and Projection (UMAP) and then clustered through the Hierarchical Density-Based Spatial Clustering of Applications with Noise (HDBSCAN) algorithm. For the number of behavioral clusters, a 0.5% minimal cluster size with run rage of 0.5%–1% was manually modified, based on the minimal temporal bout, to avoid over-clustering or under-clustering. Using the identified clusters, B-SOiD predicted behavioral groups in new DLC labeling datasets with a Random Forest classifier.

### Principal component analysis (PCA)

PCA was computed using the R function prcomp() and was analyzed further using the syndRomics R package (*23*). To determine significant components, a non-parametric permutation-based test for variance accounted for (VAF) was performed on the original 63 components with 10,000 permutations. Loadings of PC1 and PC2 variables were calculated using a non-parametric permutation test for loadings with 1,000 permutations.

### Entropy

The entropy of BGs was calculated using Shannon’s entropy as a sum across non-zero usage BGs for each mouse. Entropy = -Sum over non-zero BGs using (P(BG=BG)^*^log2(P(BG=BG)). The entropy for each individual mouse during preictal, MS, and GS was calculated and considered as repeated measures.

### Knee joint angle analysis

The coordinate of the bilateral hip, knee, and ankle (caudal foot) joints were extracted from DLC labeling. Knee joint angles were calculated using the hip, knee, and ankle labels. For each frame, vectors were defined between the hip and knee (proximal vector) and between the knee and ankle (distal vector). The knee angle between these vectors was computed using the dot product and normalized by the magnitudes of the vectors. Angular velocity was derived from the frame-to-frame differences in knee joint angles, normalized by the frame rate of the video. This allowed for quantifying the rate of change of knee joint angles over time, providing insights into the direction and speed of joint movement.

### Frequent pattern mining and association rules analysis

We used the Apriori algorithm (*59*) for frequent pattern mining with high frequency of ≥ 70% overall support and ≥ 50% for each group to ensure they serve as strong indicators of their respective classes. We then transformed the mined patterns into a “bag of patterns” representation (*60*), recorded the frequency of each pattern, and fed this representation into a Random Forest model.

### Statistics

We conducted appropriate parametric and nonparametric tests on a contingent of the Shapiro-Wilk normality test. Unless otherwise noted. Comparisons across multiple groups were made using the Friedman Test followed by Bonferroni correction for multiple comparisons. Comparisons between the two groups were made using Student’s t-test or Mann-Whitney U test. 8-day kindling behavior group usage data were transformed using a boundary adjustment for values of 0 or 1 using an adjustment of ±0.001, and then a logit transformation to normalize the data and fit a linear mixed-effects model for repeated measures. Unless otherwise noted, data were plotted in mean and standard error mean (SEM) with individual data points when applicable.

## Acknowledgments

We gratefully acknowledge Noora Rajjoub (OSU) and Miriam Najeeb (OSU) for video partition and manual Racine scoring.

## Authorship contributions

Y.Y.S: project management, conducting experiments, data analysis, and writing; J.T.: code development, data analysis, and writing; X.H.C.: computational modeling, machine learning, and writing; J.Z.: raw video processing; A.H.: raw video processing; P.Z.: conceptualization, project management, and writing; A.S.: conceptualization, project management, data processing, and writing; and B Gu: conceptualization, project management, conducting experiments, data analysis, and writing.

## Data and materials availability

All data associated with this study are present in the paper or the Supplementary Materials. All DLC and B-SOiD data and associated codes are archived on Zenodo or Github.

