## Supplemental materials for "Behavior decoding delineates seizure microfeatures and associated sudden death risks in mice"

This PDF file includes Supplemental Figures 1 to 6 and Supplemental Tables 1 and 2

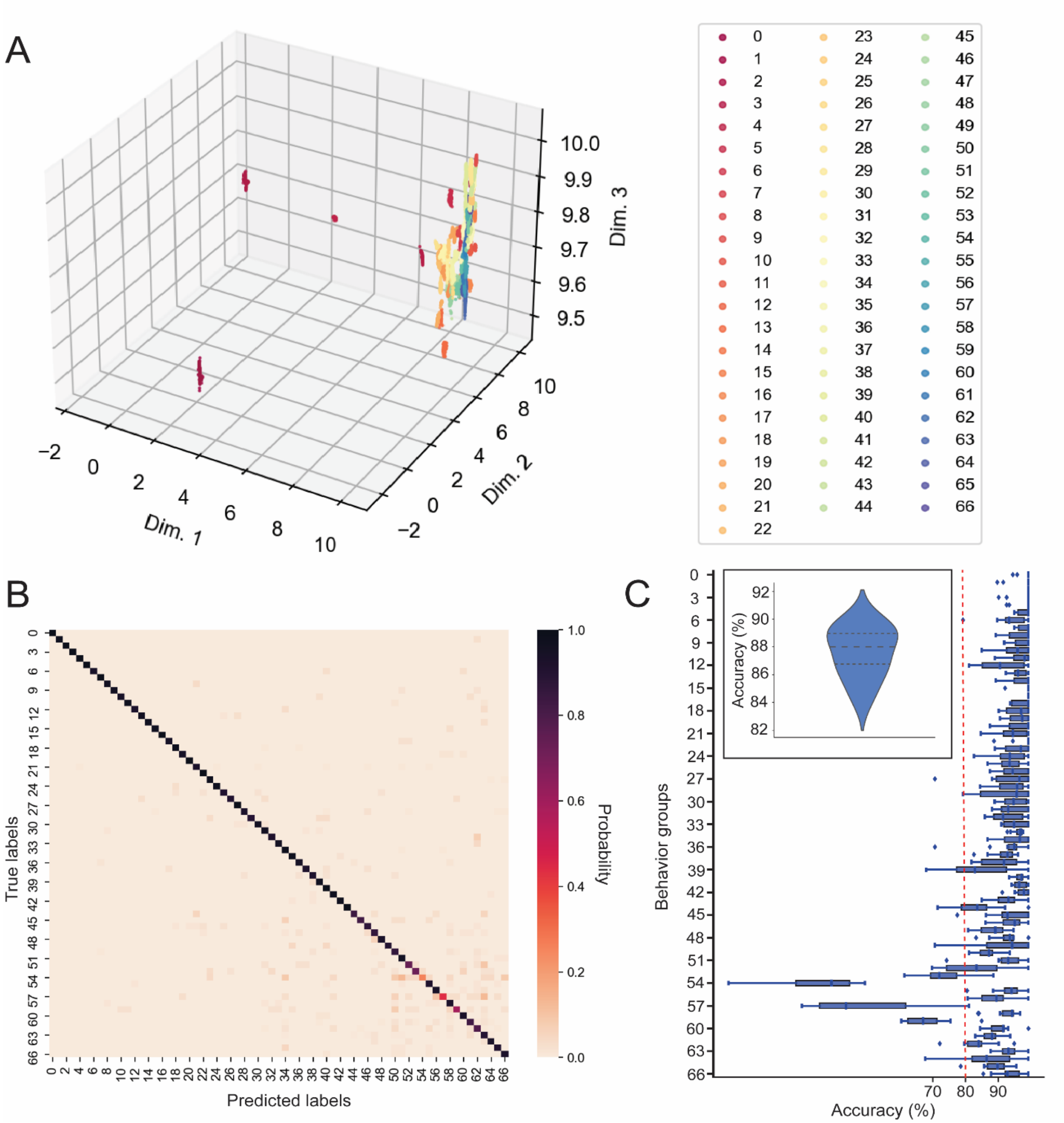


**Supplemental Figure 1**. B-SOiD model clustering and performance. (A) HDBSCAN shows the BGs clustering distribution. (B) The confusion matrix shows the classification performance of the B-SOiD model on 20% of the data. (C) The box-and-whisker plot illustrates the 10-fold cross-validation of individual BG. The outliers are included as the individual points beyond the whisker. 63 out of 67 BGs reach a median accuracy ≥ 80% except BG#53, BG#54, BG#57, and BG#59. The inset depicts the overall performance of the random forest classifier on 20% of the data with a median accuracy of 88%.

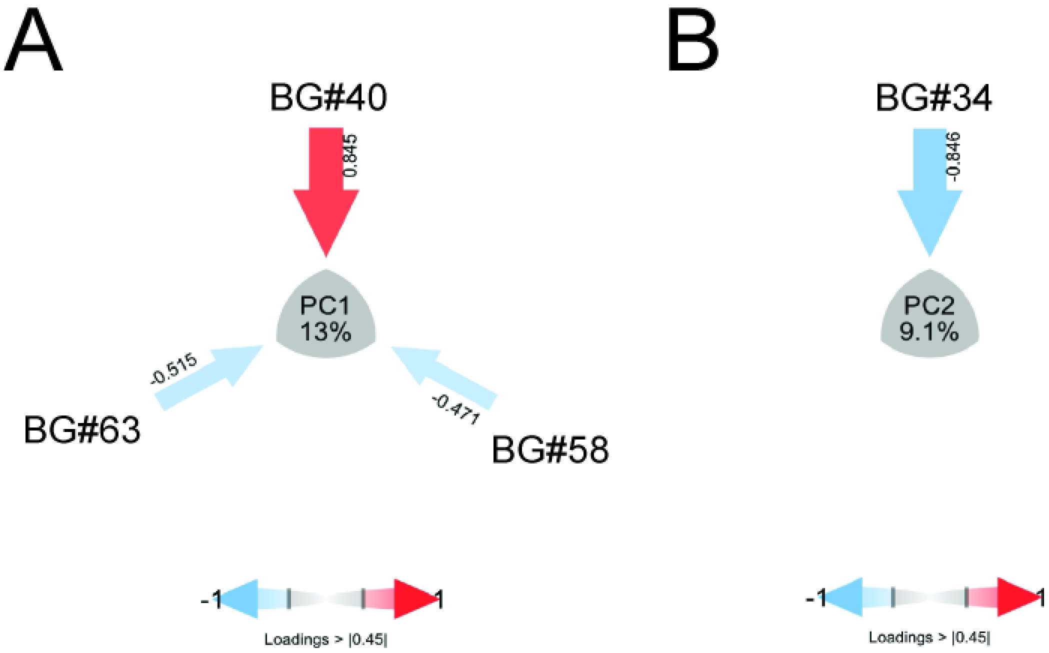


**Supplemental Figure 2**. Variables loading of PC1 and PC2. Plots of variables |loadings| > 0.45 for (A) PC1 and (B) PC2. Arrows pointing at the center of the plot represent the magnitude (arrow thickness and color saturation) and direction (color) of the loadings of selected variables.

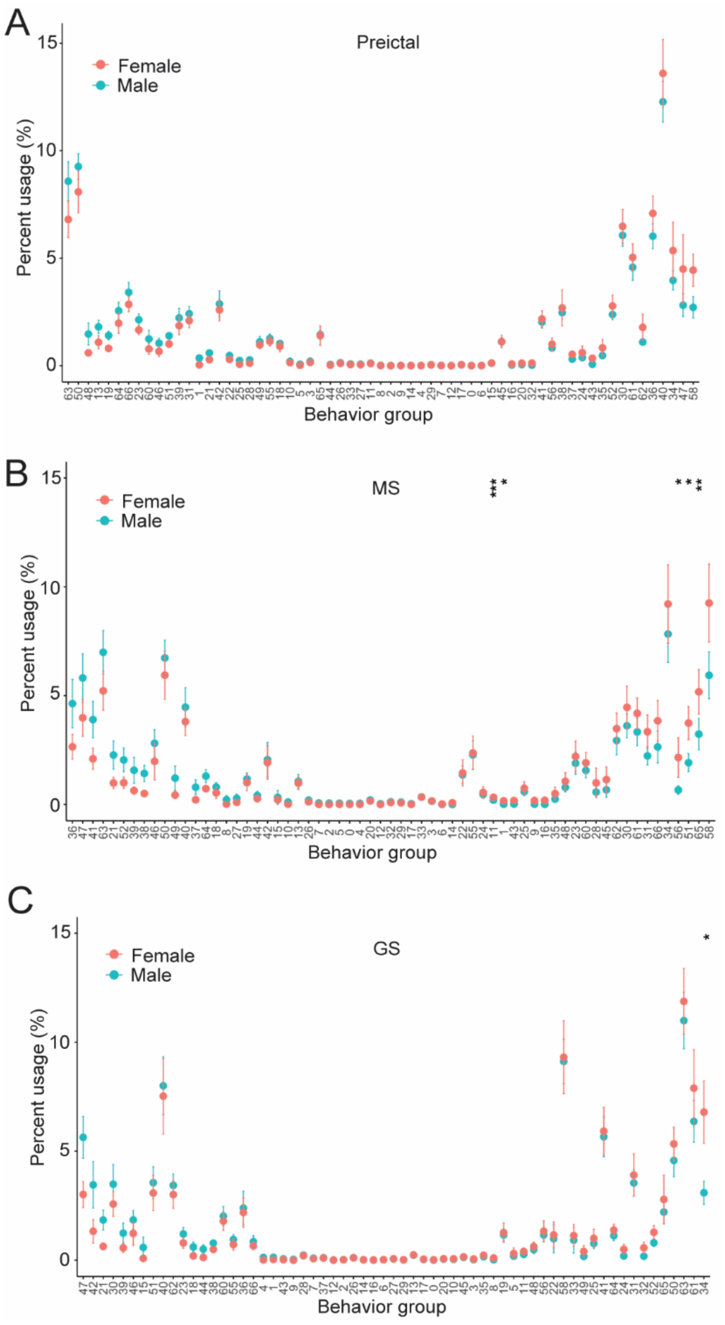


**Supplemental Figure 3**. Sex effects on behavior during preictal, MS, and GS. Percent BG usage of females (n = 72) and males (n = 147) during (A) preictal, (B) MS, and (C) GS. BGs are ranked by delta (males - females) in descending order. Data are presented as mean ± SEM. Data are analyzed using the Mann-Whitney U test, *P < 0.05, **P < 0.01, and ***P < 0.001.

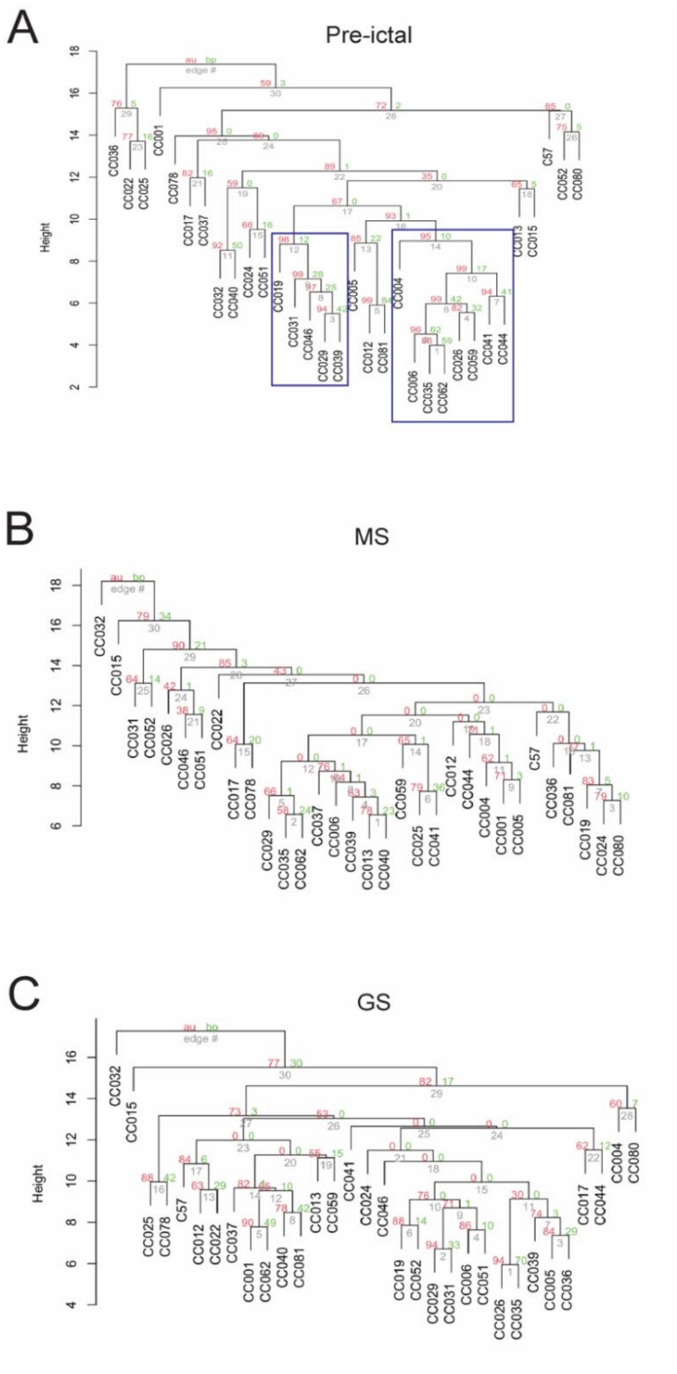


**Supplemental Figure 4**. Statistics of hierarchical clustering of mouse strains by BGs usage. Cluster dendrogram with Euclidean distance and statistics during (A) preictal, (B) MS, and (C) GS. The red number denotes approximately unbiased (AU) probability values (P values), the green number denotes bootstrap probability, and the grey number denotes cluster labels. Clusters with AU P value ≥ 95% and bootstrap > 10 are highlighted by blue rectangles and are considered to be strongly supported by data.

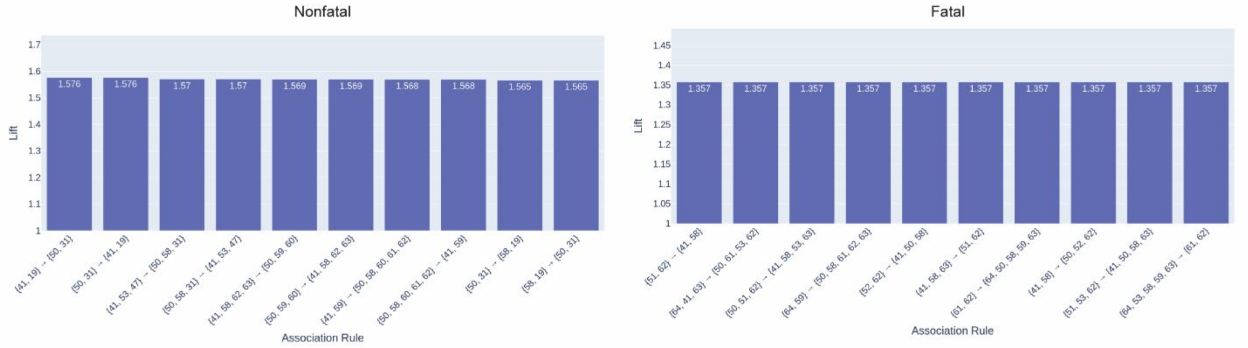


**Supplemental Figure 5**. Association rules analysis by measuring the lift of distinct collections of strongly associated itemsets for nonfatal and fatal seizures. We identified the top ten most useful itemset associations for each class.

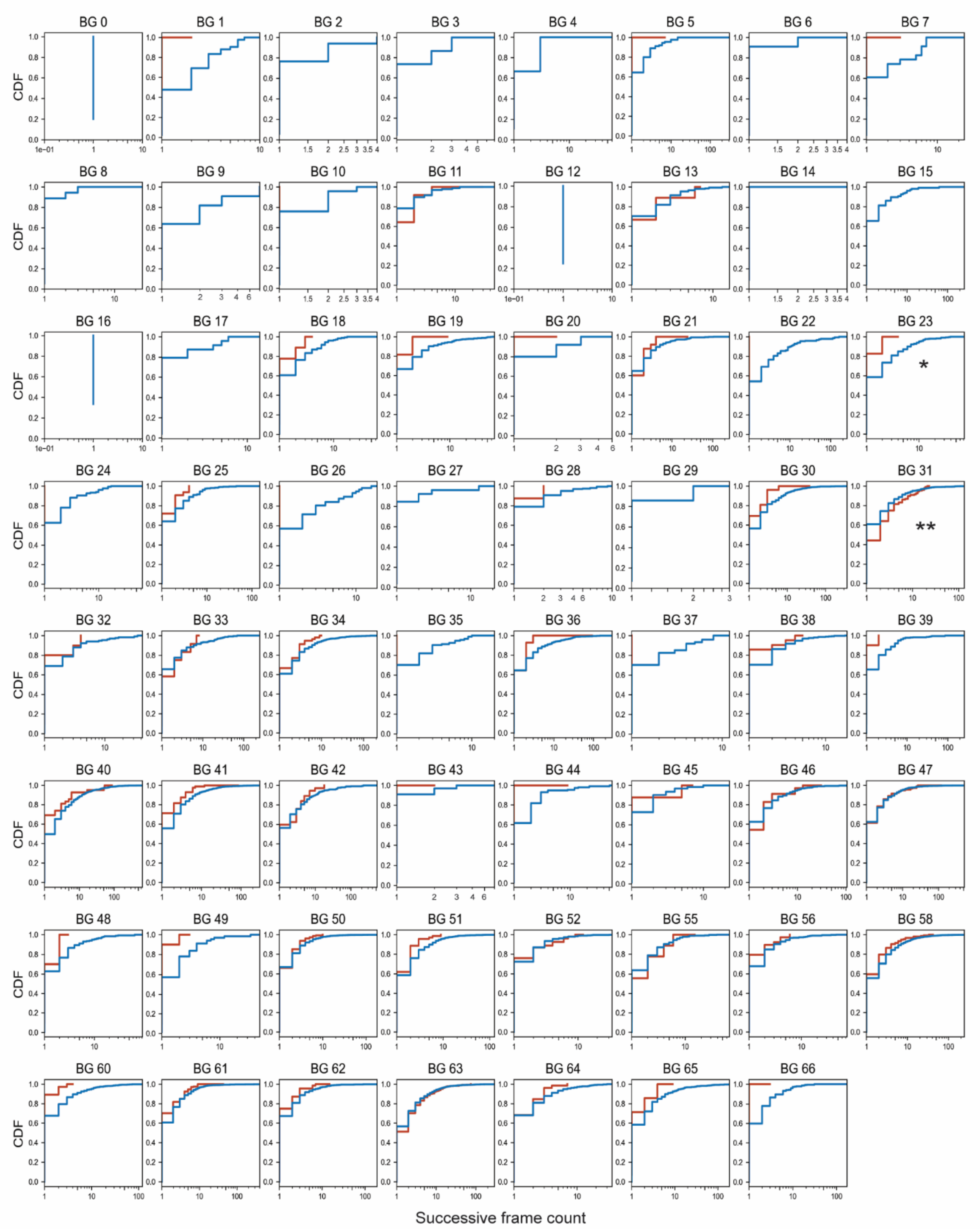


**Supplemental Figure 6**. CDF of bout duration of BGs during GS comparing fatal (n =19) to nonfatal (n = 200) seizures. CDF was analyzed using Wasserstein distance and *$P < 0.05$, **$P < 0.01$ by two-tailed Kolmogorov-Smirnov test. The X-axis is successive frames in the log scale.

**Supplemental Table 1**. Summary of dataset 1

| Strain | Males | Females | Coat color |
|---|---|---|---|
| CC001/Unc | 6 | 2 | Black |
| CC004/TauUnc | 1 | 0 | Agouti |
| CC005/TauUnc | 2 | 2 | Agouti |
| CC006/TauUnc | 3 | 6 | Agouti |
| CC012/GeniUnc | 5 | 0 | Agouti |
| CC013/GeniUnc | 6 | 0 | Agouti |
| CC015/Unc | 2 | 0 | Black |
| CC017/Unc | 3 | 7 | Agouti |
| CC019/TauUnc | 3 | 5 | Albino |
| CC022/GeniUnc | 3 | 0 | Black |
| CC024/GeniUnc | 6 | 0 | Albino |
| CC025/GeniUnc | 2 | 0 | Albino |
| CC026/GeniUnc | 2 | 8 | Agouti |
| CC029/Unc | 6 | 5 | Albino |
| CC031/GeniUnc | 9 | 6 | Albino |
| CC032/GeniUnc | 6 | 0 | Agouti |
| CC035/Unc | 3 | 6 | Agouti |
| CC036/Unc | 6 | 0 | Albino |
| CC037/TauUnc | 5 | 8 | Agouti |
| CC039/Unc | 6 | 0 | Albino |
| CC040/TauUnc | 6 | 0 | Agouti |
| CC041/TauUnc | 5 | 0 | Agouti |
| CC044/Unc | 9 | 6 | Agouti |
| CC046/Unc | 6 | 0 | Albino |
| CC051/TauUnc | 6 | 0 | Albino |
| CC052/GeniUnc | 5 | 0 | Agouti |
| CC059/TauUnc | 2 | 6 | Agouti |
| CC062/Unc | 3 | 5 | Agouti |
| CC078/TauUnc | 6 | 0 | Agouti |
| CC080/TauUnc | 6 | 0 | Albino |
| CC081/Unc | 6 | 0 | Agouti |

**Supplemental Table 2**. Behavior group ethogram

| BG# | Ethogram | BG# | Ethogram |
|---|---|---|---|
| 0 | rising | 38 | sitting up |
| 1 | standing | 39 | exploring with head bobbing |
| 2 | rearing with straub tail | 40 | scrambling up |
| 3 | minor head diving | 41 | jerking and breathing heavily |
| 4 | crouching and rearing | 42 | climbing down |
| 5 | retreating | 43 | resting and sniffing |
| 6 | head turning left | 44 | clonus |
| 7 | hunching | 45 | face grooming and turning right |
| 8 | shrinking back | 46 | clonus with swimming behavior |
| 9 | straining with head bobbing | 47 | freezing with minor head shaking |
| 10 | splaying and leaning forward | 48 | head poking |
| 11 | stretching forward | 49 | head swinging |
| 12 | head lowering | 50 | dropping |
| 13 | stepping forward | 51 | spasm |
| 14 | freezing with head down | 52 | head-turning and grooming |
| 15 | sitting with straub tail | 55 | sitting up with head lifting |
| 16 | body turning left | 56 | walking |
| 17 | crouching with head bobbing | 58 | diving |
| 18 | walking and sniffing | 60 | rearing and jerking |
| 19 | body turning right | 61 | resting and gasping |
| 20 | sniffing | 62 | clonus with straub tail |
| 21 | falling or rolling | 63 | balancing |
| 22 | stretching with straub tail | 64 | jumping |
| 23 | pushing backaward | 65 | balancing with clonus |
| 24 | hunching with head bobbing | 66 | rearing |
| 25 | stretching and head lifting | | |
| 26 | resting | | |
| 27 | righting reflexing | | |
| 28 | slow crawling | | |
| 29 | head turning right | | |
| 30 | turning left and sniffing | | |
| 31 | leaping | | |
| 32 | freezing with straub tail | | |
| 33 | tail dancing | | |
| 34 | leaning forward and sniffing | | |
| 35 | turning right and sniffing | | |
| 36 | freezing with forelimb stretching | | |
| 37 | head lifting | | |
